# Granular hydrogels improve myogenic invasion and repair after volumetric muscle loss

**DOI:** 10.1101/2023.09.28.560056

**Authors:** Gabrielle I. Tanner, Leia Schiltz, Marxa L. Figueiredo, Taimoor H. Qazi

**Author notes:** corresponding author: Taimoor H. Qazi, 206 S. Martin Jischke Drive, West Lafayette IN 47907. indicates equal contribution as first authors (Gabrielle I. Tanner, Leia Schiltz).

## Abstract

Skeletal muscle injuries including volumetric muscle loss (VML) are marked by excessive scarring and functional disability that inherent regenerative mechanisms are unable to reverse. Despite high prevalence in civilian and military populations, there is currently no effective treatment for VML but bioengineering interventions such as biomaterials that fill the VML defect to support tissue growth and repair are a promising strategy. However, traditional biomaterials developed for this purpose are rigid, non-porous constructs that hinder cell infiltration. In the present study, we test the effects of granular hydrogels on muscle repair - hypothesizing that their inherent porosity will support the invasion of native myogenic cells and their flowability will permit conformable filling of the defect site, leading to effective muscle repair. We used photocurable hyaluronic acid crosslinked with matched muscle stiffness to prepare small or large particle fragments via extrusion fragmentation and facile size sorting. In assembled granular hydrogels, particle size and degree of packing significantly influenced pore features including porosity, pore size, and pore density, as well as rheological behavior including storage moduli and yield strain. We tested the ability of granular hydrogels to support early-stage (satellite cell invasion) and late-stage (myofiber invasion) muscle repair compared to bulk hydrogels in a VML injury model in the tibialis anterior (TA) muscles of 12-14 week old mice. Histological evaluation revealed granular hydrogels supported these regenerative processes while control bulk hydrogels restricted them to the gel-tissue interface in line with the absence of invading cells. Together, these results highlight the promising potential of injectable and porous granular hydrogels in supporting endogenous repair after severe muscle injury.

## Introduction

Skeletal muscles have a robust inherent ability to regenerate after minor injuries but undergo degenerative remodeling and functional decline when severe injuries overwhelm native regenerative mechanisms. An example of this is when volumetric muscle loss (VML) leads to necrosis of a large volume of viable muscle tissue that is replaced by fibrotic scar resulting in permanent muscle weakness and painful disability. VML injuries are prevalent in both military and civilian populations and are caused for example by traumatic injury, gunshot wounds and surgical resections (1). The current clinical gold standard for treatment includes autologous grafting where viable vascularized and innervated muscle tissue is surgically transferred to the injury site (2). However, autologous grafting can lead to donor site morbidity and may be challenging to perform in older patients or those with pre-existing comorbidities (3).

As an alternative to autografts, decellularized extracellular matrices (dECMs) have been tested extensively for VML repair as they retain the bioactive composition (proteins, growth factors, RNA, etc.) of viable tissue grafts but have been cleared of DNA that could potentially elicit a foreign body immune response (4). Importantly, dECMs are acellular implants that are considered medical devices by the food and drug administration (FDA) and thus face a less stringent regulatory process than cell-based therapies (5). Implantation of dECMs have been shown to support de novo muscle fiber formation with functional recovery in animal models as well as human clinical trials (6,7). While some reports have suggested that the functional recovery after dECM implantation may not be due to muscle regeneration but could instead be attributed to fibrotic scar biomechanics (8), others have shown that dECM biomaterials can indirectly promote a pro-healing environment by local immune modulation (9). Irrespective of mechanism of action, some inherent drawbacks of dECM implants include their batch-to-batch variability, lack of control over biophysical and biochemical properties, and low porosity (10). The non-porous structure is a particularly limiting feature as interconnected porosity is needed to support cell and vessel invasion leading to tissue repair (11). Traditional methods to prepare dECM implants lead to rigid and non-porous structures that require trimming and stacking to fill the defect site which further densifies pores and hinders cell growth (12). A recent study reported on increasing the porosity of dECM implants to support myoblast proliferation by incorporating rapidly dissolvable microspheres into a dECM hydrogel (13). However, these and other similarly crosslinked scaffolds require careful handling during surgery and trimming or debridement to fit into the defect site.

As an alternative to naturally derived dECMs, acellular synthetic hydrogels with tailorable properties have been tested in VML models. These hydrogels are usually crosslinked and shaped into a bulk gel form that is implanted at the VML defect site and sutured or glued in place to enhance retention (14,15). While many studies have reported the implantation of bulk degradable hydrogels into VML defects, insights on cell migration into these hydrogels are severely lacking for example because the hydrogels degrade too quickly and are not detectable on histological follow-up (16,17). Some formulations of bulk hydrogels rely on cell-mediated degradation of the matrix to permit invasion. However, this outside-in mode of cell migration usually restricts regenerative activity at the hydrogel-viable tissue interface and can miss the time window to revive a necrotic tissue core in large VML defects (18). Biomaterials that support rapid cell invasion while providing a supporting scaffolding are highly desirable in a VML context to establish bioactivity and create a regenerative microenvironment for tissue repair.

Granular hydrogels are an emerging class of biomaterials that support rapid cell infiltration due to their inherent microporous structure (19,20). Granular hydrogels are assembled through the packing of microscale particle building blocks. Interstitial pores naturally exist between adjacent particles and this porosity can be further enhanced by modifying particle size (21), shape (22), or degree of packing (23). A unique advantage of granular hydrogels is that porosity can be decoupled from particle mechanics or degradability, as a result allowing the fabrication of highly modular hydrogels with multifunctional properties (24). Granular hydrogels are also injectable through clinically relevant syringe needles and catheters as particles can flow on the application of force and rapidly recover on force removal (25,26). This allows minimally invasive delivery to sites of tissue damage and extrusion into volumetric defects with smooth filling and contouring of uneven defect edges such as those found in clinical scenarios (27,28). Although granular hydrogels have been applied to treat brain stroke (29), myocardial infarction (30), peripheral nerve defects (31), and dermal wounds (32), their application to treat VML defects has not been explored so far.

In this study, we evaluated cell invasion and tissue repair after implantation of bulk and granular hydrogels in a mouse model of VML injury. A photocurable polymer was used to fabricate granular hydrogels using fragmented particles. We report the inclusion of a facile size-sorting step after particle fabrication that results in small or large particles that, when packed, form granular hydrogels with significantly different pore features and rheological behavior. In a mouse tibialis anterior (TA) VML model, we observe a regenerative myogenic response characterized by the invasion of activated satellite cells and newly regenerated fibers into porous granular hydrogels compared to almost non-existent invasion in traditional bulk hydrogels. Our findings demonstrate the advantages of granular hydrogels over traditional bulk hydrogels in supporting skeletal muscle regeneration.

## Results and Discussion

### Particle fragmentation and granular hydrogel formation

Norbornene-modified hyaluronic acid (Nor-HA) hydrogels were used in this study owing to their demonstrated biocompatibility, tunable mechanical properties, and fast crosslinking chemistry (Fig. 1a) (33). Bulk hydrogel properties were tailored to identify formulations that mimicked the elastic modulus of native skeletal muscle (∼11-13 kPa) by varying polymer concentrations and degrees of crosslinking. A 2% w/v Nor-HA formulation with 50% degree of crosslinking (corresponding to 4 mM DTT) had stiffness values of ∼12 kPa and was used throughout the rest of this study (Fig. 1b). Hydrogel microparticles were obtained using an extrusion fragmentation technique where bulk hydrogel is extruded through successively smaller syringe needles, as reported earlier (23,25,34). This technique is quick and results in a high particle yield compared to microfluidics-based approaches that additionally involve excessive washing steps to remove oil and surfactant (35). Fluorescence microscopy images showed polygonal-shaped particles that became smaller in size with each step of the fragmentation process (Fig. 1c). Quantification of particle size by intensity thresholding and particle analysis in FIJI software showed that particles obtained after passing through 30G needles were significantly smaller than particles obtained after passing through 18G, 21G, and 23G needles (Fig. 1d). The relatively high variability in size is inherent to this method of particle fabrication as deformation and fracture of polygonal shaped particles during extrusion is a random process (25). Thus, granular hydrogel formulations containing fragmented particles have a mix of small and large particles, which can make it challenging to study the impact of particle size. To overcome this issue, we introduced a facile sorting step after fragmentation to obtain small and large particle fractions from unsorted batches (Fig. 1e). Standard 100 μm cell strainers fitted with nylon meshes retained larger particles while allowing smaller ones to pass through (Fig. 1f). Fluorescence microscopy (Fig. 1g) and quantitative particle size analysis confirmed significant differences between the two sorted fractions (Fig. 1h). Despite the effectiveness of sorting, some small particles were still observed in the large particle fraction likely because filtration is highly dependent on particle orientation at the time of contact with the mesh as also observed in other studies (36). Using custom-built meshes to obtain targeted particle size is a potential alternative to post-fragmentation sorting, as reported by the Zenobi-Wong lab that used metal sieves (37,38), although this can sometimes lead to elongated particle strands instead of isotropic particles (39).

**Figure 1:**
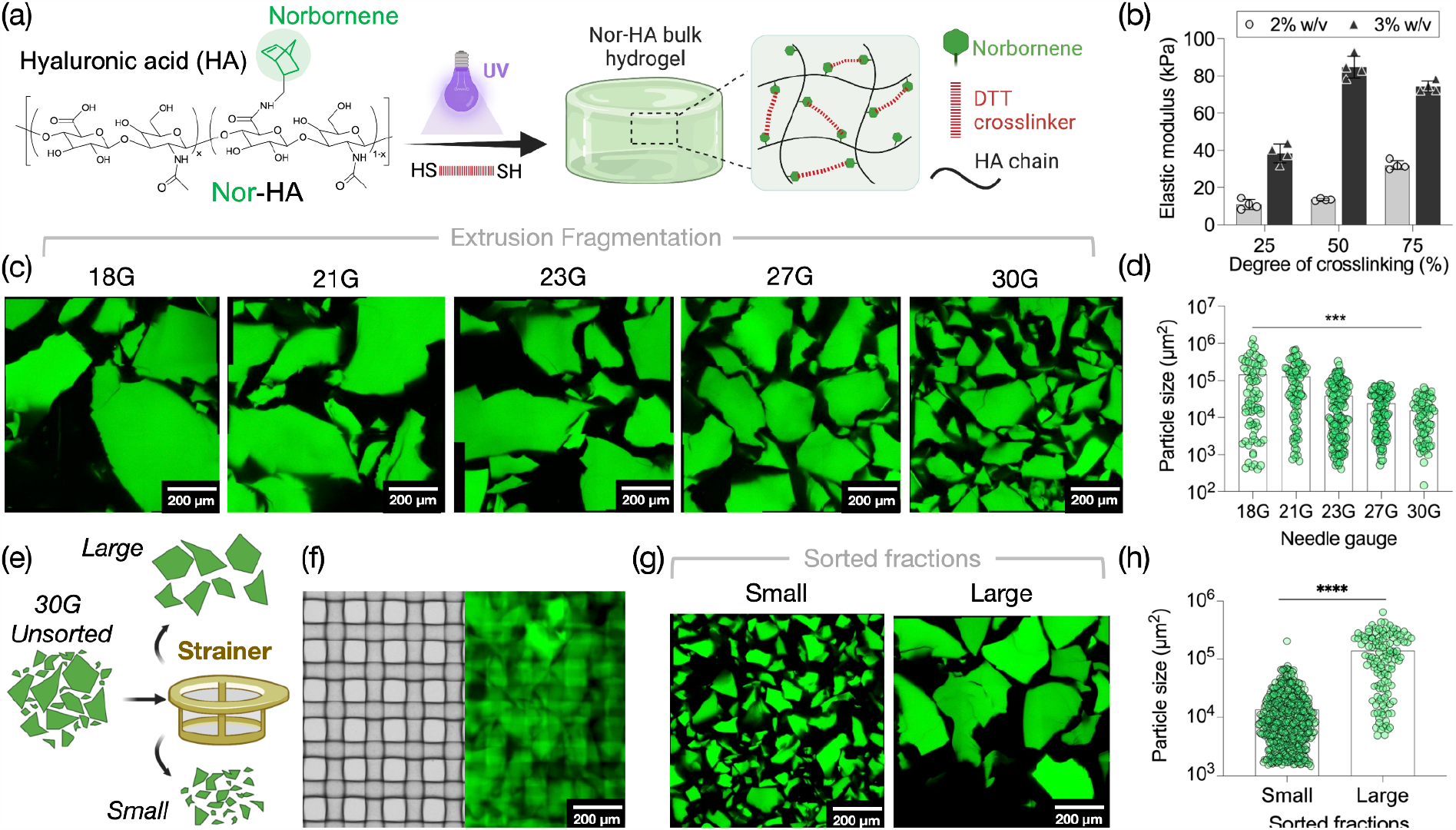
Fabrication and sorting of fragmented hydrogel particles. (a) Schematic showing the covalent crosslinking of norbornene-modified hyaluronic acid (Nor-HA) bulk hydrogels. (b) The elastic modulus of hydrogels is tuned by varying the polymer concentration or degree of crosslinking. (c) Morphology and (d) size of particles obtained at every step of the extrusion fragmentation process after passing through syringe needles of increasing gauge. (e) Use of a cell strainer to sort fragmented particles into small and large fractions. (f) Strainer mesh with an average pore size of 100 μm retains large particles. (g) Morphology and (h) size of small and large particles obtained after sorting. Scale bars: 200 μm.

### Granular hydrogel porosity and pore features

Granular hydrogels with interstitial pores are formed when particles are packed together via forced aggregation (40,41). These pores ultimately support cell invasion into granular hydrogels in a tissue repair context (42,43). Prior work has shown that particle size, shape, and the degree of particle packing can influence interstitial pore features including pore size, number, orientation, and the overall hydrogel void fraction (21–23). Here, we investigated how sorting fragmented particles into small and large fractions influences hydrogel porosity compared to standard unsorted fractions. To do this, unsorted and sorted fragmented particle fractions were packed via vacuum filtration ensuring a very high degree of packing; these groups were referred to as the “high packing” groups. To increase porosity, the “high packing” groups were diluted with the equivalent of 15% and 30% v/v PBS, resulting in “medium packing” and “low packing” granular hydrogel groups, respectively. Compared to centrifugation-based differences in degrees of packing (21), this method of unjamming highly aggregated particles offers greater control over hydrogel porosity and properties (44). Pores were examined using confocal microscopy to image fluorescent FITC-dextran molecules that were infiltrated through the granular hydrogel (Fig. 2a). In hydrogels made with unsorted particles, both large and small particles were observed with smaller particles often appearing to plug pores between adjacent large particles. This was confirmed by the coefficient of variation of pore size which was the highest for unsorted particle fractions compared to sorted fractions at all degrees of packing. Void fraction defined as the fraction of pores per region of interest and averaged across multiple regions of interest ranged from ∼5% to ∼20% across groups (Fig. 2b). As expected, unjamming the high packed granular hydrogels by dilution with PBS increased the void fraction for all groups. Notably, granular hydrogels made with small particles had the highest void fraction whereas the ones made with unsorted particles had the lowest void fraction independent of degree of packing (Fig. 2b). This is in contrast to results reported with spherical particles where void fraction is correlated with particle size (21,45), highlighting the substantial influence of particle shape on structural properties of granular hydrogels. The number of pores per region of interest was also highest for granular hydrogels made with small particles, averaging, for example, ∼90 pores/ROI compared to ∼30 and ∼25 for the unsorted and large particle groups at the highest degree of packing (Fig. 2c). Granular hydrogels made with small particles had the lowest average pore size, in line with observations made using spherical particles (21).

**Figure 2:**
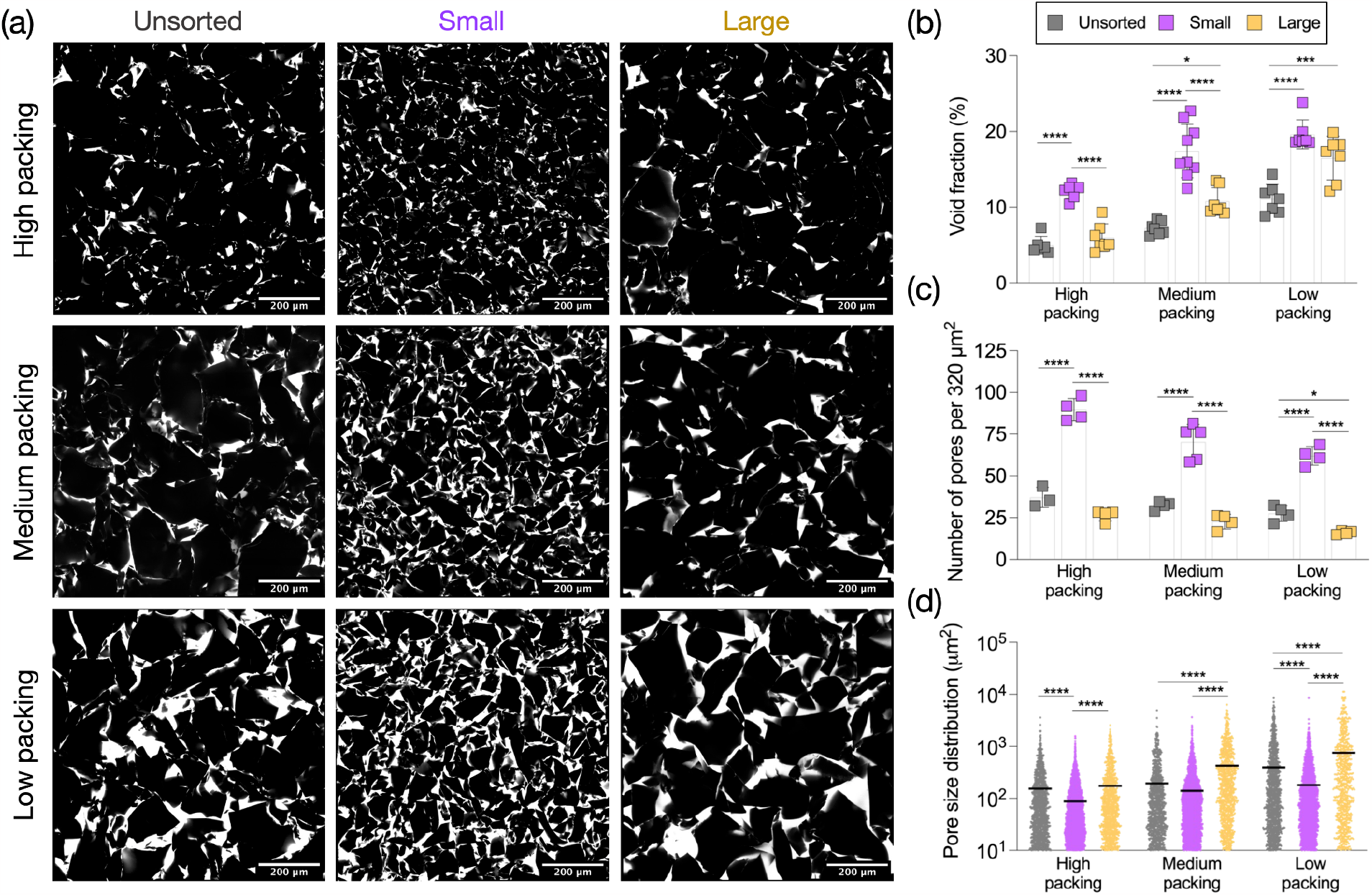
Fragmented particle size and packing impact pore features of granular hydrogels. (a) Representative confocal microscopy images of interstitial pores (white) between fragmented particles (black) showing differences in pore morphology with changes in particle size and degree of packing. Scale bars: 200 μm. (b) The average void fraction of granular hydrogels is highest with small particle fragments compared to unsorted and large fragment groups. (c) The average number of pores per region of interest is highest with small particle fragments compared to unsorted and large fragment groups at all degrees of packing. (d) The average pore size is lowest with small particle fragments compared to unsorted and large fragment groups at all degrees of packing.

### Granular hydrogel injectability and rheological behavior

Granular hydrogels show strain-yielding and self-healing behavior that allows particles to flow past one another on the application of force and undergo rapid recovery when the force is removed (46). These features have enabled the application of granular hydrogels as extrudable filaments or host slurries in 3D printing, and as injectable biomaterials that can be introduced at sites of tissue damage minimally invasively using standard syringe needles and catheters. Traditional bulk hydrogels with dynamic crosslinks display similar shear thinning and self-healing properties but lack the interparticle porosity that is a distinguishing feature of injectable granular hydrogels. To study how fragmented particle size and packing influenced the rheological behavior of granular hydrogels, we used oscillatory shear rheology on unsorted, small, and large particle fragments and with low, medium, and high degrees of packing. Strain sweeps performed over a range of 0.5% to 100% strain demonstrated the characteristic strain-yielding behavior of granular hydrogels. Notably, these plots illustrate the gradual decrease in the storage moduli (G’) and increase in the loss moduli (G”) with increasing strain values, ultimately reaching the yield strain where G” overtakes G’ indicating a switch from primarily solid-like to liquid-like behavior (Fig. 3a). As anticipated, the different degrees of packing result in hydrogels with different storage moduli and yield strains, and thus the strain sweep plots start at different G’ values and the curves for G” and G’ cross-over at different strain values, but all tested groups show a generally similar behavior. Hydrogels were subjected to periodically applied regimes of low (0.5%) and high (500%) shear strain to evaluate mechanical recovery after yielding (Fig. 3b). All granular hydrogel groups tested showed a near-complete recovery of their storage moduli values after repetitively being subjected to high strains. In general, storage moduli (G’) for all granular hydrogel groups were equal to or above ∼1000 Pa, in line with prior observations (25,47). This is in contrast to granular hydrogels made with spherical particles that have comparably lower G’ values even at higher degrees of packing (21). These differences can be attributed to the particle shape (e.g., smooth spheres vs. rough polygonal fragments) that can impact inter-particle friction, sliding, and interlocking when granular hydrogels are subjected to shear, leading to differences in moduli and yielding behavior. Between different groups, the storage moduli decreased with lower degrees of packing (high packing > medium packing > low packing), and granular hydrogels made with the large particle fraction had comparatively the highest storage moduli (Fig. 3c). The yield strain values were comparable between the groups, ranging from ∼7% to ∼20%, and in the range of expected values as per prior studies by us and others.

**Figure 3:**
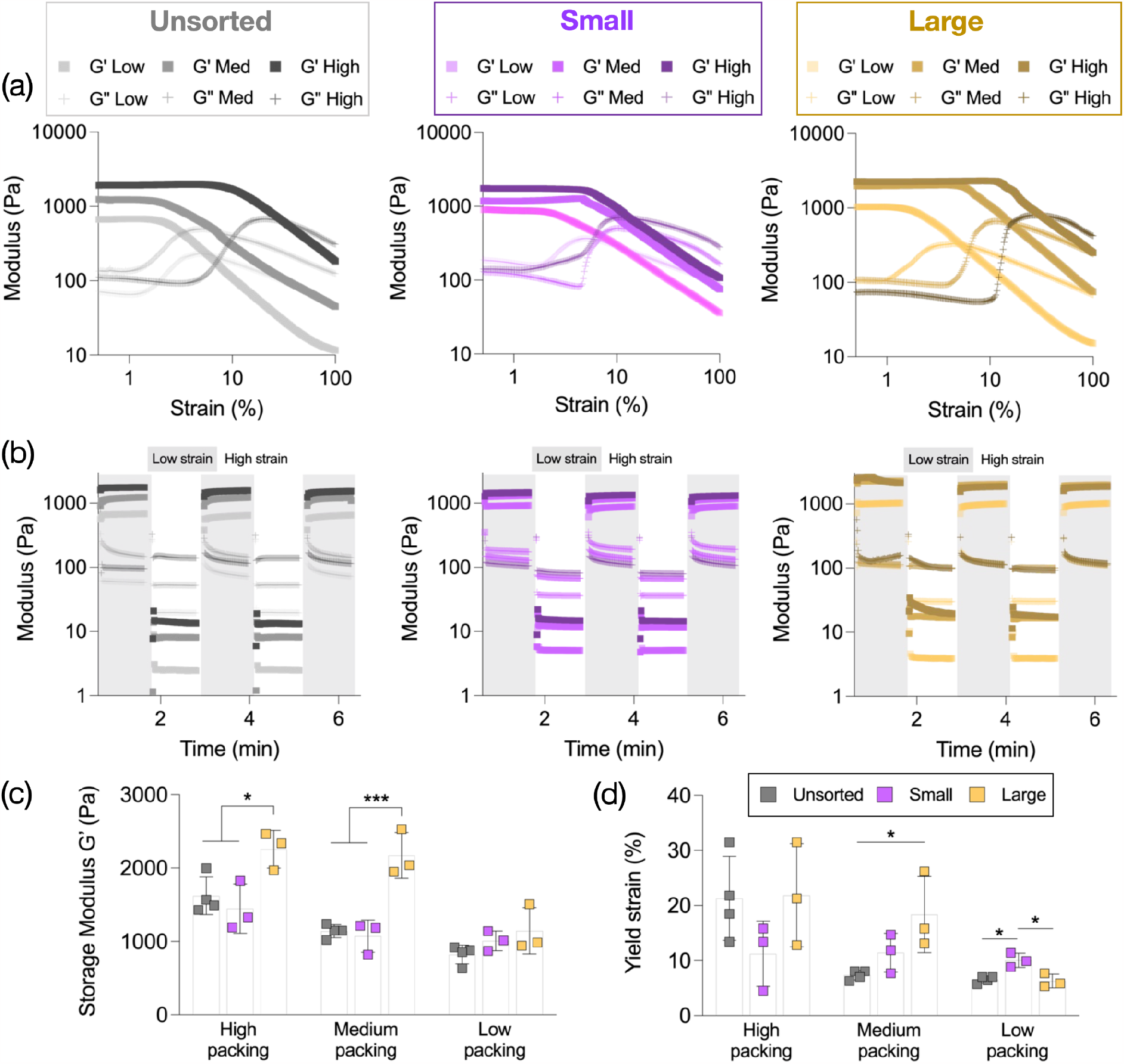
Rheological behavior of granular hydrogels made with unsorted and sorted particle fractions. (a) Representative strain sweeps (0.5-100%) of granular hydrogels made with unsorted (left), small (middle), and large (right) particle fractions with low, medium, or high degrees of packing. (b) Representative time sweeps of granular hydrogels showing strain-yielding and self-healing behavior after being repetitively subjected to successive periods of low (0.5%) and high (500%) strains. (c) Average storage moduli (G’ derived at 0.5% strain) and (d) average yield strain (% strain at crossover point when G”>G’) of granular hydrogels.

### Application of granular hydrogels in VML injury model

We created full-thickness VML defects in mouse tibialis anterior (TA) muscles using a 2.5 mm biopsy punch to remove a significant portion of viable muscle tissue. The *in vivo* study design included implantation of either bulk or granular (small or large particle fractions at medium packing) hydrogels immediately after injury with animal euthanasia and histological evaluation at 1-week and 4-week time points (Fig. 4a). The goal of this *in vivo* study was to investigate the early and late endogenous cellular response to biomaterial implantation, in particular comparing the effects of particle size in the granular hydrogels’ ability to support myogenic cell invasion and establishing the benefits of this class of biomaterials compared to traditionally used bulk hydrogels. The TA is an easily accessible muscle that has previously been used in VML models (48,49); its mid-belly area is wide enough to permit the creation of a 2.5 mm wide full-thickness defect (Fig. 4b). This defect size corresponds to ∼25-30% loss of viable tissue, satisfying the definition of a critical VML injury (50). Importantly, when VML defect samples over a total of 28 TA muscles were weighed to compare variability in defect size, there were no significant differences observed across the three hydrogel groups (Fig. 4c).

**Figure 4:**
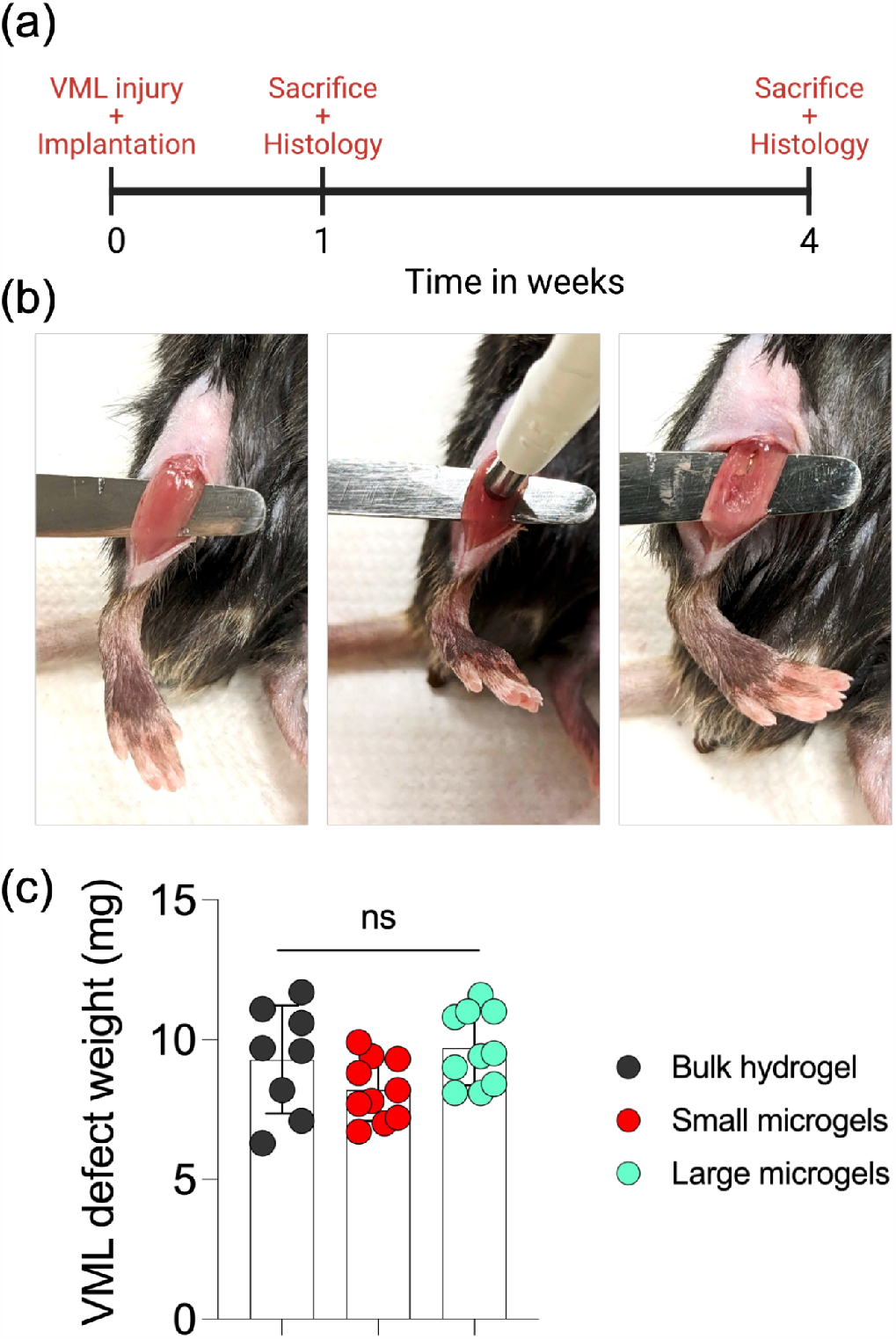
Mouse tibialis anterior volumetric muscle loss model. (a) Study timeline with endpoints at 1 week and 4 weeks after initial injury. (b) Images showing the creation of the VML defect. (c) Biopsy weight showing consistency in creating VML defects in 28 TA muscles randomly assigned across three hydrogel implantation groups.

### Endogenous cell invasion into hydrogels after VML injury

At study endpoints (1 week and 4 weeks), mice were euthanized and TA muscles were cryopreserved for histological evaluation. We opted to create longitudinal sections with myofibers running along the length of the section as this orientation allows improved visualization of the VML defect and the hydrogel. After 1 week, the injury area in the TA mid-belly was clearly identifiable by the absence of F-Actin (magenta) staining and the clustering of nuclei (cyan) indicating an endogenous cellular response. Bulk or granular hydrogels (yellow) were also visible inside the defect region (Fig. 5a). In comparison, normal myofiber and nuclei morphology was observed in the sham (positive control) group. In TAs that received granular hydrogels, a small fraction of particles were visible outside of the defect region suggesting particle transport in the absence of inter-particle crosslinks and likely enhanced due to animal movement and muscle contraction. Cell invasion into hydrogels was qualitatively evaluated at a higher magnification and it showed a complete lack of cell infiltration within bulk hydrogels and substantial infiltration within the interstitial pores of granular hydrogels (Fig. 5b). Quantification of nuclei density within ROIs marked by the hydrogel showed that granular hydrogels made with small particles had the highest cell invasion of ∼12%, those made with large particles had cell invasion of ∼7%, whereas the bulk hydrogels had ∼0% invasion (Fig. 5c). The bulk hydrogels were not expected to support cell invasion as they lacked any degradable crosslinks. In comparison, granular hydrogels can support cell invasion due to interstitial porosity even when particles are non-degradable. This is an especially attractive feature of granular hydrogels as their ability to support rapid cell infiltration throughout the gel structure prevents the formation of a necrotic core which forms with traditional bulk degradable hydrogels that support a slow mode of outside-in migration (18). At 4 weeks, the bulk and granular hydrogels were still clearly visible within the VML defect region (Fig. 5d) with higher magnification images showing cells restricted to the outside periphery of bulk hydrogels and invading porous granular hydrogels (Fig. 5e). Quantification at 4 weeks revealed there were no significant differences in cell invasion between granular hydrogels made with large and small particles and that cell density had reduced to levels comparable to the sham group (Fig 5f). Bulk hydrogels still had the lowest cell density of ∼0%. Overall, these data highlight the superior ability of granular hydrogels to support endogenous cell invasion after a volumetric muscle injury in comparison to traditional bulk hydrogels. We previously reported that granular hydrogels injected subcutaneously are invaded by multiple cell types including macrophages, myofibroblasts, and endothelial cells and the cellularity data in Fig. 5 likely reflects this cellular diversity (51). Here, we next investigated the invasion of muscle-specific cells focusing on the early and late-stage myogenic repair response.

**Figure 5:**
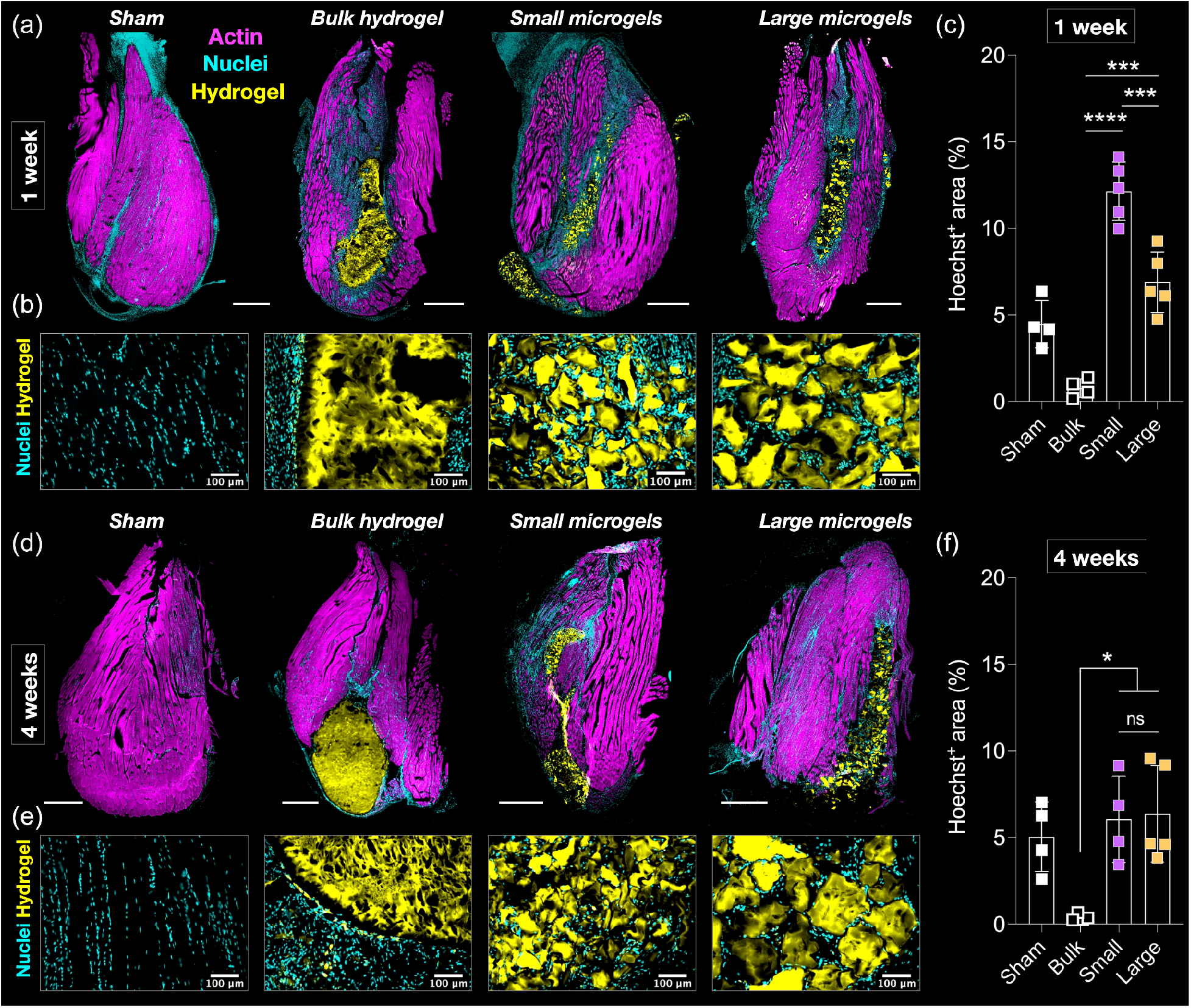
Hydrogel morphology and endogenous cell invasion in mouse VML defects. Representative tile-scan images of longitudinal muscle sections at the (a) 1-week and (d) 4-week time points showing the retention of bulk or granular hydrogels in the VML injury regions compared to uninjured sham controls. Scale bars: 1000 μm. High magnification images at the (b) 1-week and (e) 4- week time points showing endogenous cells located around the outside surface of bulk hydrogels and successfully invading porous granular hydrogels made with small or large particle fractions. Scale bars: 100 μm. Quantification of cell invasion into hydrogels at the (c) 1-week and (f) 4-week time points.

### Myogenic repair after hydrogel implantation in VML injury model

The regenerative response to muscle injury is marked by the activation of Paired box protein 7 (Pax7+) quiescent satellite cells, proliferation of myogenic progenitors, and differentiation into myofibers [ref]. As regeneration progresses with the maturation and eventual fusion of myogenitor cells, the expression of myogenin (MyoG) and myoblast determination protein 1 (MyoD) increases whereas Pax7 expression decreases (52). To evaluate how hydrogel implantation supported this repair process, we first stained muscle cryosections with Pax7+ antibody. Pax7+ myogenic cells including activated satellite cells and myoblasts were readily observed in and around the VML defect region at 1 and 4 week timepoints (Fig. 6a). As expected, Pax7+ cells were restricted to the outside periphery of bulk hydrogels whereas they had infiltrated the porous granular hydrogels. In comparison, the sham group had a very minimal number of Pax7+ cells, in line with reports that quiescent satellite cells represent less than 1% of the total cell population in homeostatic tissue in the absence of injury-related signals. Strikingly, Pax7+ cells infiltrating the granular hydrogels had an elongated morphology that is suggestive of an activated myogenic progenitor phenotype (e.g., Pax7+MyoD-MyoG- or Pax7+MyoD+MyoG-) although recent reports have highlighted the morphological diversity of quiescent satellite cells that often have numerous protrusions extending along the surface of the myofiber niche (53,54). Quantification of Pax7+ invasion showed comparable levels between the two granular hydrogel groups and a higher density at 1 week compared to 4 weeks (Fig. 6c). This is in line with the expected trajectory of myogenic regenerative response where satellite cell activation peaks in the early stages after injury and declines thereafter. To evaluate differentiation of myogenic progenitors into new myofibers, we next stained muscle sections with embryonic myosin heavy chain (eMHC) antibody that marks newly regenerated myofibers (55). As expected, eMHC+ myofibers were not observed in the sham group but were readily observed in other groups around the injury region at 1 and 4 week timepoints (Fig. 6b). Interestingly, multinucleated eMHC+ myofibers were observed adjacent to bulk hydrogels indicating a regenerative response that is limited to the outer periphery of bulk non-porous hydrogels. In contrast, long and continuous eMHC+ myofibers had invaded porous granular hydrogels. A robust invasion of eMHC+ myofibers with wide and continuous multinucleated structures was evident in granular hydrogels made with large particles, whereas relatively discontinuous structures was observed in granular hydrogels made with small particles. This could be due to differences in inter-particle pore size, connectivity, and tortuosity. Quantitative evaluation of eMHC+ invasion did not reveal a significant difference in signal intensity per ROI between the small and large particle groups (Fig. 6d). Importantly, eMHC+ area increased, albeit non-significantly, between weeks 1 and 4 indicating persistent myofiber regeneration.

**Figure 6:**
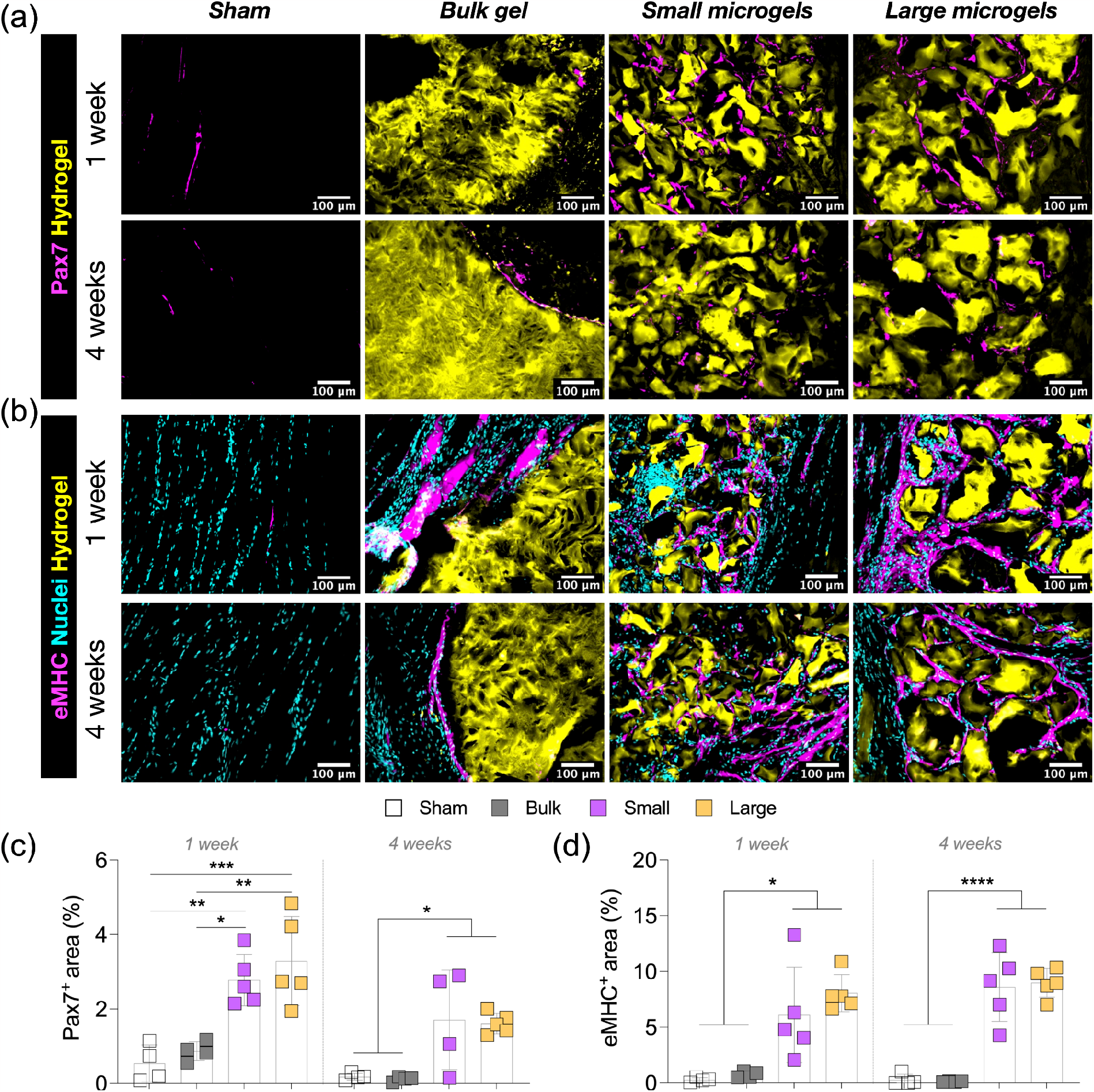
Myogenic progenitor cell and myofiber invasion into hydrogels. (a) Representative images showing (a) Pax7+ cells (magenta) and (b) eMHC+ cells and myofibers (magenta) in and around the vicinity of the implanted hydrogels (yellow) at the 1-week and 4-week time points. Scale bars: 100 μm. Quantification of (c) Pax7+ cell invasion and (d) eMHC+ cell and myofiber invasion into the hydrogels.

## Conclusions

Both granular hydrogel formulations tested *in vivo* (made with either small or large fragmented particles) supported the rapid invasion of cells including Pax7+ satellite cells and activated myogenic progenitors and multinucleated eMHC+ myofibers. In stark contrast, bulk hydrogels with similar mechanical properties prevented any cells from invading. *In vitro* characterization showed granular hydrogel porosity and pore features can be tailored by modifying particle size and degree of packing that could potentially improve cell invasion. With their modular design and highly customizable design parameters, granular hydrogels are promising biomaterials to treat muscle injuries such as volumetric muscle loss.

## Experimental section

### Granular hydrogel formation

Nor-HA was synthesized as previously described (33). Briefly, HA modified with tetrabutylammonium salt (HA-TBA) was dissolved in anhydrous dimethyl sulfoxide (DMSO). Dimethyl aminopyridine, norbornene-2-carboxylic acid, and di-tert-butyl dicarbonate were added to the mixture and allowed to react for 20 hours at 45°C. The reaction was quenched with an equal volume of cold water, and the Nor-HA product was then dialyzed for 10 days in DI water, filtered, frozen, lyophilized until dry, and stored at −20°C until further use. ^1^H NMR analysis was used to determine the degree of modification.

Nor-HA polymer was dissolved in phosphate buffered saline (PBS), mixed with LAP photoinitiator (1 mg.mL^-1^) and DTT crosslinker (4 mM), and transferred to a 3 mL syringe prior to crosslinking with UV at 20 mW.cm^2^ for 2 minutes. Bulk hydrogels were then extruded through syringe needles with subsequently smaller inner diameters starting with 18G, 21G, 23G, 27G, and 30G. Fragmented hydrogel microparticles obtained were cleaned by washing three times with excess PBS. A cell strainer with a 100 μm mesh size was used to sort fragmented particles into small and large particle fractions. Granular hydrogels were formed through particle packing by vacuum filtration. Briefly, particles were suspended in PBS and transferred to a Durapore membrane filter placed on a Buchner funnel attached to a vacuum line. The aggregated particles collected directly from the membrane were referred to as the “high packing” granular hydrogel group. The “medium packing” group was obtained by diluting these granular hydrogels with 15% v/v PBS, and the “low packing” group was obtained by diluting with 30% v/v PBS.

### Mechanical characterization

Compression testing of bulk hydrogels was carried out on an Anton Paar MCR 702e dynamic mechanical analyzer to determine elastic moduli. Briefly, cylindrical hydrogel samples were crosslinked under UV light (Omnicure S1500) at an intensity of 20 mW.cm^-2^ for 1 minute inside PDMS molds. The samples were washed with PBS and allowed to swell overnight before unconfined uniaxial compression testing up to 20% strain at a strain rate of 1 mm.min^-1^. The elastic modulus value for each sample was calculated using the slope of the stress-strain curve in the 5%-15% strain region.

Oscillatory shear rheology with a parallel plate geometry and a gap height of 1 mm was carried out on an Anton Paar MCR 702e dynamic mechanical analyzer, as previously described. Strain sweeps were carried out from 0.5% to 500% strain at 1 Hz to determine yield strains (defined as the strain at which G” becomes higher than G’) and characterize strain-yielding behavior. Time sweeps were carried out with cyclical application of low (0.5%) and high (500%) strains to characterize self-healing and mechanical recovery behavior. Storage and loss moduli were determined at 0.5% strain.

### Porosity characterization

Particles were carefully suspended in PBS containing 2-5 mg.mL^-1^ FITC-dextran (1 M.Da) and packed via vacuum filtration. Additional FITC-dextran solution was added if preparing medium or low packing samples. Hydrogels were transferred using a spatula into custom-made PDMS molds adhered on a glass slide and covered on top with a 1.5 glass coverslip. Care was taken to avoid bubbles. An upright Zeiss LSM 800 confocal microscope was used to obtain volumetric z-stacks in multiple regions of interest within the hydrogel and across at least n=3 samples. In the images, the particles appeared dark as they were unlabeled and the interstitial pores between particles appeared bright indicating efficient FITC-dextran infiltration and retention. Confocal stacks were processed using FIJI software. Briefly, after binarization and thresholding, the analyze particles function was used to quantify pore size, number of pores, and void fraction.

### Animal surgery and biomaterial implantation

All procedures involving animals were approved by the Purdue Institutional Animal Care and Use Committee (PACUC protocol #03230023680). 10-14 week-old male C57BL/6 mice were used for this study. Mice were anesthetized with isoflurane and hindlimbs were shaved and treated with depilatory cream. Analgesics were administered subcutaenously once prior to surgery and every 24 hours for three days after surgery. Bilateral full thickness volumetric muscle loss (VML) injuries were created in the tibialis anterior (TA) muscles of mice using a 2.5 mm sterile biopsy punch. For this, an incision was made laterally to the TA using a sterile scalpel, and blunt dissection was performed to identify and separate the TA from the skin and loosen the fascia. A flat spatula was placed underneath the TA and the biopsy punch was applied to the mid-belly portion of the TA in a twisting motion until a full thickness defect was achieved. The VML sample was collected and weighed. For the sham group (positive control) skin incision and blunt dissection of the TA from skin was performed but no VML injury was created. A total of four groups were tested with TA muscles across animals randomly assigned to: (1) Sham, (2) Bulk hydrogel, (3) Granular hydrogel containing small particles, (4) Granular hydrogel containing large particles. All hydrogels for *in vivo* application were fluorescently labeled with FITC-dextran. For the bulk hydrogel group, the VML defect area was dried by application of sterile gauze and 20 μL of the polymer precursor solution was introduced into the VML defect and photocrosslinked for 30 seconds. For the granular hydrogel groups, 20 μL of the hydrogel was extruded directly into the VML defect using a syringe attached to a 23G needle. Following implantation, the skin was carefully sutured and the mice were returned to their cages for recovery.

### Histological analyses

At study endpoints (1 week and 4 weeks after VML injury), animals were anesthetized using isoflurane and euthanized by cervical dislocation. TA muscles were harvested from all animals and fixed in 4% PFA for 24 hours, followed by immersion in 15% and 30% sucrose for 24 hours. Muscle samples were then embedded in OCT-Tissue Tek and cryopreserved using liquid nitrogen cooled isopentane. Tissues were sectioned in the longitudinal direction using a cryostat at a section thickness of 15 μm and adhered onto Superfrost slides. Sections were stained with Hoechst (PI62249; Fisher Scientific) and either Phalloidin (A22287; Alexa Fluor-647), Pax7 (DSHB), or embryonic myosin heavy chain (DSHB), followed by corresponding fluorescent secondary antibodies. Sections were covered with Prolong gold antifade mounting medium and dried overnight. Imaging was performed with a 10x (tile scans) or 20x objective using a Keyence BZ-X810 inverted fluorescence microscope and processed using FIJI software. Quantification of cell invasion was performed by first outlining the region of interest (ROI) containing the biomaterial (identified by FITC signal) and then determining total Hoechst, Pax7, or eMHC intensity within that ROI.

### Statistical analyses

Data were analyzed on GraphPad Prism software. In vitro experiments were repeated at least three independent times. The Shapiro-Wilk test was used to determine normal distribution of data points. Differences between two groups were tested with two-tailed student’s t-test. Differences between more than two groups were tested with One-Way ANOVA with Bonferroni’s post-hoc test. Levels of significance were set at ^*^p<0.05, ^**^p<0.01, ^***^p<0.001, ^****^p<0.0001.

## Acknowledgments

The authors gratefully acknowledge new faculty start-up funds from the Weldon School of Biomedical Engineering (to T.H.Q.), and financial support towards the Summer Undergraduate Research Fellowship (SURF) program from the Purdue University College of Engineering and the Engineering Undergraduate Research Office (EURO) (to G.I.T. and L.S.). The authors thank William Schoenlein and the Purdue BME Vet Tech team for assistance with animal studies, John Harwood and the Purdue Interdepartmental NMR Facility for assistance with polymer analysis, Xiaoguang Zhu and the Purdue Imaging Facility for the use of confocal microscopes, Linlin Li and the Umulis lab for use of the cryostat, and Hamood ur Rehman and the Markworth lab for assistance with H&E staining.

## References

1. Testa S, Fornetti E, Fuoco C, Sanchez-Riera C, Rizzo F, Ciccotti M, et al. The war after war: Volumetric muscle loss incidence, implication, current therapies and emerging reconstructive strategies, a comprehensive review. Biomedicines. 2021;9(5).

2. Hurtgen BJ, Ward CL, Leopold Wager CM, Garg K, Goldman SM, Henderson BEP, et al. Autologous minced muscle grafts improve endogenous fracture healing and muscle strength after musculoskeletal trauma. Physiol Rep. 2017;5(14):1–16.

3. Smoak MM, Mikos AG. Advances in biomaterials for skeletal muscle engineering and obstacles still to overcome. Mater Today Bio [Internet]. 2020;7(May):100069. Available from: 10.1016/j.mtbio.2020.100069

4. Nuge T, Liu Z, Liu X, Ang BC, Andriyana A, Metselaar HSC, et al. Recent advances in scaffolding from natural-based polymers for volumetric muscle injury. Molecules. 2021;26(3).

5. Christman KL. Regenerative medicine: Biomaterials for tissue repair. Science (80-). 2019;363(6425):340–1.

6. Sicari BM, Peter Rubin J, Dearth CL, Wolf MT, Ambrosio F, Boninger M, et al. An acellular biologic scaffold promotes skeletal muscle formation in mice and humans with volumetric muscle loss. Sci Transl Med. 2014;6(234).

7. Dziki J, Badylak S, Yabroudi M, Sicari B, Ambrosio F, Stearns K, et al. An acellular biologic scaffold treatment for volumetric muscle loss: results of a 13-patient cohort study. npj Regen Med [Internet]. 2016;1(1):1–12. Available from: 10.1038/npjregenmed.2016.8

8. Greising SM, Rivera JC, Goldman SM, Watts A, Aguilar CA, Corona BT. Unwavering Pathobiology of Volumetric Muscle Loss Injury. Sci Rep [Internet]. 2017;7(1):1–14. Available from: 10.1038/s41598-017-13306-2

9. Sadtler K, Powell JD, Wolf MT, Elisseeff JH, Estrellas K, Pardoll DM, et al. Developing a pro-regenerative biomaterial scaffold microenvironment requires T helper 2 cells. Science (80-). 2016;352(6283):366–70.

10. Zhang X, Chen X, Hong H, Hu R, Liu J, Liu C. Decellularized extracellular matrix scaffolds: Recent trends and emerging strategies in tissue engineering. Bioact Mater [Internet]. 2022;10(September 2021):15–31. Available from: 10.1016/j.bioactmat.2021.09.014

11. Annabi N, Nichol JW, Zhong X, Ji C, Koshy S, Khademhosseini A, et al. Controlling the porosity and microarchitecture of hydrogels for tissue engineering. Tissue Eng - Part B Rev. 2010;16(4):371–83.

12. Hodde J, Hiles M. Constructive soft tissue remodelling with a biologic extracellular matrix graft: Overview and review of the clinical literature. Acta Chir Belg. 2007;107(6):641–7.

13. Eugenis I, Wu D, Hu C, Chiang G, Huang NF, Rando TA. Scalable macroporous hydrogels enhance stem cell treatment of volumetric muscle loss. Biomaterials. 2022;290(September).

14. Narayanan N, Jia Z, Kim KH, Kuang L, Lengemann P, Shafer G, et al. Biomimetic glycosaminoglycan-based scaffolds improve skeletal muscle regeneration in a Murine volumetric muscle loss model. Bioact Mater [Internet]. 2021;6(4):1201–13. Available from: 10.1016/j.bioactmat.2020.10.012

15. Ziemkiewicz N, Hilliard GM, Dunn AJ, Madsen J, Haas G, Au J, et al. Laminin-111-Enriched Fibrin Hydrogels Enhance Functional Muscle Regeneration Following Trauma. Tissue Eng - Part A. 2022;28(7–8):297–311.

16. Carleton MM, Locke M, Sefton M V. Methacrylic acid-based hydrogels enhance skeletal muscle regeneration after volumetric muscle loss in mice. Biomaterials [Internet]. 2021;275(May):120909. Available from: 10.1016/j.biomaterials.2021.120909

17. Dienes J, Browne S, Farjun B, Amaral Passipieri J, Mintz EL, Killian G, et al. Semisynthetic Hyaluronic Acid-Based Hydrogel Promotes Recovery of the Injured Tibialis Anterior Skeletal Muscle Form and Function. ACS Biomater Sci Eng. 2021;7(4):1587–99.

18. Basurto IM, Passipieri JA, Gardner GM, Smith KK, Amacher AR, Hansrisuk AI, et al. Photoreactive Hydrogel Stiffness Influences Volumetric Muscle Loss Repair. Tissue Eng - Part A. 2022;28(7–8):312–29.

19. Riley L, Schirmer L, Segura T. Granular hydrogels: emergent properties of jammed hydrogel microparticles and their applications in tissue repair and regeneration. Curr Opin Biotechnol [Internet]. 2019;60:1–8. Available from: 10.1016/j.copbio.2018.11.001

20. Griffin DR, Weaver WM, Scumpia PO, Di Carlo D, Segura T. Accelerated wound healing by injectable microporous gel scaffolds assembled from annealed building blocks. Nat Mater. 2015;14(7):737–44.

21. Qazi TH, Muir VG, Burdick JA. Methods to Characterize Granular Hydrogel Rheological Properties, Porosity, and Cell Invasion. ACS Biomater Sci Eng. 2022;8(4):1427–42.

22. Qazi TH, Wu J, Muir VG, Weintraub S, Gullbrand SE, Lee D, et al. Anisotropic Rod-Shaped Particles Influence Injectable Granular Hydrogel Properties and Cell Invasion. Adv Mater. 2022;34(12):1–12.

23. Muir VG, Qazi TH, Shan J, Groll J, Burdick JA. Influence of Microgel Fabrication Technique on Granular Hydrogel Properties. ACS Biomater Sci Eng. 2021;7(9):4269–81.

24. Mealy JE, Chung JJ, Jeong HH, Issadore D, Lee D, Atluri P, et al. Injectable Granular Hydrogels with Multifunctional Properties for Biomedical Applications. Adv Mater. 2018;30(20).

25. Muir VG, Qazi TH, Weintraub S, Torres Maldonado BO, Arratia PE, Burdick JA. Sticking Together: Injectable Granular Hydrogels with Increased Functionality via Dynamic Covalent Inter-Particle Crosslinking. Small. 2022;18(36).

26. Highley CB, Song KH, Daly AC, Burdick JA. Jammed Microgel Inks for 3D Printing Applications. Adv Sci. 2019;6(1).

27. Nih LR, Sideris E, Carmichael ST, Segura T. Injection of Microporous Annealing Particle (MAP) Hydrogels in the Stroke Cavity Reduces Gliosis and Inflammation and Promotes NPC Migration to the Lesion. Adv Mater. 2017;29(32):1–8.

28. Xin S, Deo KA, Dai J, Pandian NKR, Chimene D, Moebius RM, et al. Generalizing hydrogel microparticles into a new class of bioinks for extrusion bioprinting. Sci Adv. 2021;7(42):1–12.

29. Wilson KL, Pérez SCL, Naffaa MM, Kelly SH, Segura T. Stoichiometric Post-Modification of Hydrogel Microparticles Dictates Neural Stem Cell Fate in Microporous Annealed Particle Scaffolds. Adv Mater. 2022;34(33):1–12.

30. de Rutte JM, Koh J, Di Carlo D. Scalable High-Throughput Production of Modular Microgels for In Situ Assembly of Microporous Tissue Scaffolds. Adv Funct Mater. 2019;29(25):1–10.

31. Hsu RS, Chen PY, Fang JH, Chen YY, Chang CW, Lu YJ, et al. Adaptable Microporous Hydrogels of Propagating NGF-Gradient by Injectable Building Blocks for Accelerated Axonal Outgrowth. Adv Sci. 2019;6(16).

32. Pruett LJ, Jenkins CH, Singh NS, Catallo KJ, Griffin DR. Heparin Microislands in Microporous Annealed Particle Scaffolds for Accelerated Diabetic Wound Healing. Adv Funct Mater. 2021;31(35).

33. Gramlich WM, Kim IL, Burdick JA. Synthesis and orthogonal photopatterning of hyaluronic acid hydrogels with thiol-norbornene chemistry. Biomaterials [Internet]. 2013;34(38):9803–11. Available from: 10.1016/j.biomaterials.2013.08.089

34. Kessel B, Lee M, Bonato A, Tinguely Y, Tosoratti E, Zenobi-Wong M. 3D Bioprinting of Macroporous Materials Based on Entangled Hydrogel Microstrands. Adv Sci. 2020;7(18):1–13.

35. Krüger AJD, Bakirman O, Guerzoni LPB, Jans A, Gehlen DB, Rommel D, et al. Compartmentalized Jet Polymerization as a High-Resolution Process to Continuously Produce Anisometric Microgel Rods with Adjustable Size and Stiffness. Adv Mater. 2019;31(49).

36. Hirsch M, D’Onofrio L, Guan Q, Hughes J, Amstad E. 4D printing of Metal-Reinforced double network granular hydrogels. Chem Eng J [Internet]. 2023;473(August):145433. Available from: 10.1016/j.cej.2023.145433

37. Puiggalí-Jou A, Asadikorayem M, Maniura-Weber K, Zenobi-Wong M. Growth factor– loaded sulfated microislands in granular hydrogels promote hMSCs migration and chondrogenic differentiation. Acta Biomater. 2023;166:69–84.

38. Asadikorayem M, Surman F, Weber P, Weber D, Zenobi-Wong M. Zwitterionic Granular Hydrogel for Cartilage Tissue Engineering. Adv Healthc Mater. 2023;2301831:1–14.

39. Rizzo R, Bonato A, Chansoria P, Zenobi-Wong M. Macroporous Aligned Hydrogel Microstrands for 3D Cell Guidance. ACS Biomater Sci Eng. 2022;8(9):3871–82.

40. Widener AE, Bhatta M, Angelini TE, Phelps EA. Guest-host interlinked PEG-MAL granular hydrogels as an engineered cellular microenvironment. Biomater Sci. 2021;9(7):2480–93.

41. Xin S, Dai J, Gregory CA, Han A, Alge DL. Creating Physicochemical Gradients in Modular Microporous Annealed Particle Hydrogels via a Microfluidic Method. Adv Funct Mater. 2020;30(6):1–9.

42. Sheikhi A, de Rutte J, Haghniaz R, Akouissi O, Sohrabi A, Di Carlo D, et al. Microfluidic-enabled bottom-up hydrogels from annealable naturally-derived protein microbeads. Biomaterials. 2019;192(October 2018):560–8.

43. Daly AC, Riley L, Segura T, Burdick JA. Hydrogel microparticles for biomedical applications. Nat Rev Mater [Internet]. 2020;5(1):20–43. Available from: 10.1038/s41578-019-0148-6

44. Anderson AR, Nicklow E, Segura T. Particle fraction is a bioactive cue in granular scaffolds. Acta Biomater. 2022;150:111–27.

45. Caldwell AS, Rao V V., Golden AC, Anseth KS. Porous bio-click microgel scaffolds control hMSC interactions and promote their secretory properties. Biomaterials [Internet]. 2020;232(October 2019):119725. Available from: 10.1016/j.biomaterials.2019.119725

46. Xin S, Wyman OM, Alge DL. Assembly of PEG Microgels into Porous Cell-Instructive 3D Scaffolds via Thiol-Ene Click Chemistry. Adv Healthc Mater. 2018;7(11):1–7.

47. Hirsch M, Charlet A, Amstad E. 3D Printing of Strong and Tough Double Network Granular Hydrogels. Adv Funct Mater. 2021;31(5).

48. Dunn A, Haas G, Madsen J, Ziemkiewicz N, Au J, Johnson D, et al. Biomimetic sponges improve functional muscle recovery following composite trauma. J Orthop Res. 2022;40(5):1039–52.

49. Quarta M, Cromie M, Chacon R, Blonigan J, Garcia V, Akimenko I, et al. Bioengineered constructs combined with exercise enhance stem cell-mediated treatment of volumetric muscle loss. Nat Commun [Internet]. 2017;8:1–17. Available from: 10.1038/ncomms15613

50. Anderson SE, Han WM, Srinivasa V, Mohiuddin M, Ruehle MA, Moon JY, et al. Determination of a critical size threshold for volumetric muscle loss in the mouse quadriceps. Tissue Eng - Part C Methods. 2019;25(2):59–70.

51. Qazi TH, Wu J, Muir VG, Weintraub S, Gullbrand SE, Lee D, et al. Anisotropic Rod - Shaped Particles Influence Injectable Granular Hydrogel Properties and Cell Invasion. Adv Mater. 2021;2109194.

52. ChargÉ SBP, Rudnicki MA. Cellular and Molecular Regulation of Muscle Regeneration. Physiol Rev [Internet]. 2004 Jan 1;84(1):209–38. Available from: 10.1152/physrev.00019.2003

53. Ma N, Chen D, Lee JH, Kuri P, Hernandez EB, Kocan J, et al. Piezo1 regulates the regenerative capacity of skeletal muscles via orchestration of stem cell morphological states. Sci Adv. 2022;8(11):1–15.

54. Kann AP, Hung M, Wang W, Nguyen J, Gilbert PM, Wu Z, et al. An injury-responsive Rac-to-Rho GTPase switch drives activation of muscle stem cells through rapid cytoskeletal remodeling. Cell Stem Cell [Internet]. 2022;29(6):933-947.e6. Available from: 10.1016/j.stem.2022.04.016

55. Kotsaris G, Qazi TH, Bucher CH, Zahid H, Pöhle-Kronawitter S, Ugorets V, et al. Odd skipped-related 1 controls the pro-regenerative response of fibro-adipogenic progenitors. npj Regen Med. 2023;8(1).

